# Direct precise measurement of the stall torque of the flagellar motor in *E. coli*

**DOI:** 10.1101/2021.10.28.466275

**Authors:** Bin Wang, Guanhua Yue, Rongjing Zhang, Junhua Yuan

**Affiliations:** Hefei National Laboratory for Physical Sciences at the Microscale and Department of Physics, University of Science and Technology of China, Hefei, Anhui 230026, China

## Abstract

The flagellar motor drives the rotation of flagellar filaments, propeling the swimming of flagellated bacteria. The maximum torque the motor generates, the stall torque, is a key characteristics of the motor function. Direct measurements of the stall torque carried out three decades ago suffered from large experimental uncertainties, and subsequently there were only indirect measurements. Here, we applied magnetic tweezer to directly measure the stall torque in *E. coli*. We precisely calibrated the torsional stiffness of the magnetic tweezer, and performed motor resurrection experiments at stall, accomplishing a precise determination of the stall torque per torque-generating unit (stator unit). From our measurements, each stator passes 2 protons per step, indicating a tight coupling between motor rotation and proton flux.

---

The flagellar rotary motor in *E*.*coli* converts transmembrane proton flux into flagellar rotation, propelling the swimming of bacteria. A motor torque-generating unit (a stator unit) is composed of four MotA and two MotB proteins, forming two proton-conducting transmembrane channels. Driven by a proton electrochemical potential difference across the cytoplasmic membrane (the proton motive force, PMF), protonation and de-protonation of Asp-32 in MotB at the cytoplasmic end of either channel induce conformational changes of a stator unit, which exerts force on the periphery of the rotor via electrostatic and steric interaction^1,2^, and the resulting torque is transmitted to the flagellar filament via a series of molecular shaft composed of a rod and a flexible hook. A motor can contain up to 11 functionally independent stators, exchanging with a membrane pool of stators on a timescale of 1 min ^3-11^.

A key property of the flagellar motor is its torque-speed relationship, measuring how much torque it generates at different speed. This relationship was measured earlier with the electro-rotation method to vary the external torque ^12,13^, and subsequently by labelling different sizes of latex beads to shortened filament stub or by changing medium viscosity to vary the viscous load ^14-18^. The motor torque is maximum at stall, and maintains approximately constant up to a knee speed, after which it drops rapidly to zero. In *E. coli* at room temperature, the knee speed is about 170 Hz, and the speed at zero torque is about 300 Hz. The stall torque per stator is one of the key characteristics of the flagellar motor. As the motor is in equilibrium at stall, one can infer how many protons a stator passes per revolution from the value of the stall torque and the PMF.

The earliest direct measurement of the stall torque for a wildtype flagellar motor was performed by flowing medium to stall the tethered cell, giving a value in the range of 1000 − 5000 pN×nm due to large experimental uncertainty ^19^. A subsequent measurement was conducted with optics tweezers, resulting in a value of about 4500 pN×nm ^20^. As it was not able to determine the number of stators in a wildtype motor in those experiments, the number was usually assumed to be about 8, resulting in a value of stall torque per stator in the range of 125 − 625 pN×nm or about 563 pN×nm. Subsequently, indirect measurements were performed, by labeling 1.0-μm-diameter bead to shortened filament stub and assuming that the torque under this high load is the same as the stall torque. Those indirect measurements generated a value of the stall torque per stator in the range of 146 − 320 pN×nm ^4,15,21^. The most recent indirect measurement with a sodium-driven chimeric motor in *E. coli* resulted in an estimate of each stator passing about 37 ions per revolution, inconsistent with the value of 26 or 52 ions as each motor takes 26 steps per revolution ^22,23^. This promoted the proposal of the mechanism of loose coupling between the proton flux the motor rotation ^24^, in direct contrast to the long-holding view that the proton flux and the motor rotation are tightly coupled ^25-27^.

Here, to resolve these inconsistencies, we applied magnetic tweezer to perform motor resurrection experiments at stall, so that we can directly measure both the stall torque and the stator number, resulting in a precise determination of the stall torque per stator.

## Result

### Motor resurrection at stall

A schematic of the experimental setup is presented in Fig. 1B (see details in Materials and Methods). Two permanent magnets generate the magnetic field for the tweezer. A magnetic bead was attached to the hook of the motor. If the motor was pulled to stall by the magnetic tweezer, motor torque was balanced by externally applied torsion:

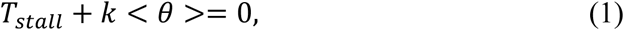

where *T*_*stall*_ was the stall toque of motors, *k* was the torsional stiffness of magnetic tweezer, < *θ* > was the angular change for the orientation of the magnetic bead, relative to that when the motor torque was zero. *k* depends on the magnitude of the magnetic field, and the number and alignment of magnetic nano-particles in individual magnetic beads ^28^. *k* was calibrated by measuring the rotational thermal fluctuations of the bead orientation < *δθ*^2^ > and applying the equipartition theorem *k* = *k*_*B*_*T*/< *δθ*^2^ >. In practice, there were apparent differences among individual beads, so it was necessary to calibrate the magnetic tweezer for each bead attaching to a de-energized motor. And the motor was then energized to generate torque.

**Fig. 1.**
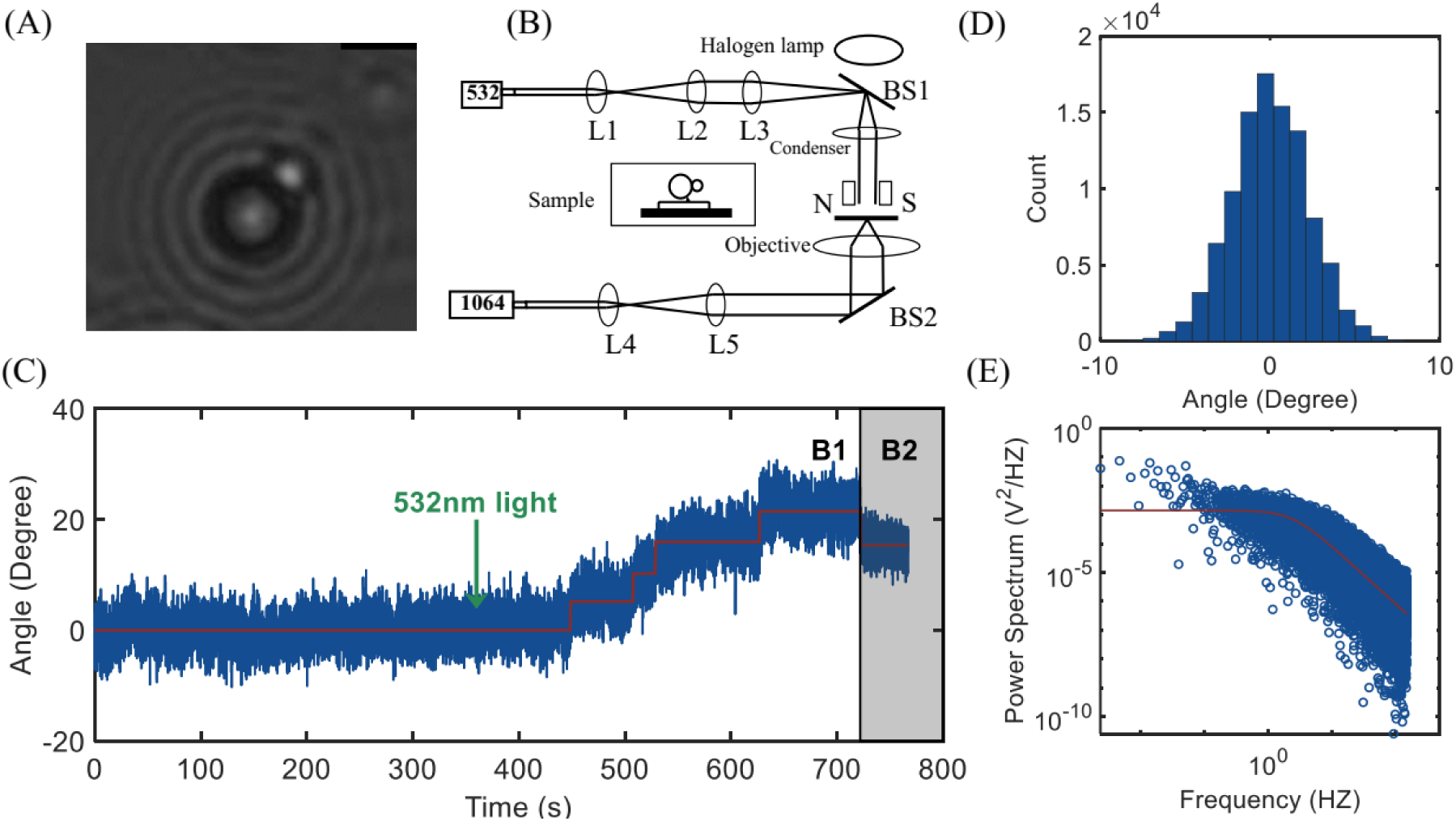
Scheme for precisely measuring the stall torque of flagellar motor. (A). An example bright-field image of the beads. (B). The experimental setup. L1, L2, L3, L4, L5 were convex lenses, and BS1 and BS2 were dichroic mirrors. Inset was a sketch of the sample and details was shown in Materials and Method. (C) An typical trace of motor resurrection at stall. The original PMF was eliminated, and the magnetic bead was undergoing Brownian motion in the magnetic tweezer. Then 532 nm light was turned on at *t* = 360 s, and the PMF was restored with the stator newly being recruited into the motor, as shown in the stepping of the bead orientation. The magnetic field was B_1_ initially, and changed quickly to B_2_ at *t* = 720 s by moving the magnets to a closer distance from the sample with a PC-controlled translational stage. (D). Histogram of the orientation trace of bead in inactive motor from (C). (E). The typical power spectrum for the orientation trace of the magnetic bead. The red line is a Lorentizian fit with the rolloff frequency extracted to be *f*_*c*_ = 2.65 Hz.

The *E. coli* K12 strain JY9, which was deleted for the genes *cheY* and *fliC* and carried mutated *flgE* expressing the hook protein with a tetracysteine motif, was transformed with the plasmid pTrc99aPR that expresses the light-driven proton pump proteorhodopsin. The hook of the motor was biotinylated, and a streptavidin-coated 2.8-μm-diameter magnetic bead was labelled to the hook in motility buffer containing 23 mM NaN_3_. Then a 1-μm-diameter biotinylated bead was manipulated with optical trap to attach to the magnetic bead as a fiducial marker for characterizing the orientation of the magnetic bead. The original PMF was eliminated by respiratory inhibition with NaN_3_ in several minutes, and the stators came off the motor on a timescale of minutes ^29,30^. So the rotor-hook-magnetic bead system would be undergoing rotational Brownian motion with magnetic restraint. To calibrate the stiffness, the motor was recorded for about 360 s in magnetic constraint of suitable strength which could be controlled by adjusting the distance between the magnets and the sample. An example bright-field image of the beads with a Brownian motion trace is shown in Fig. 1A, and the corresponding histogram of the bead orientation in Fig. 1D. The double-bead method allowed us to determine the orientation of the magnetic bead to a precision of 0.03 degree^2^ (by analyzing the angular variance of beads stuck to glass surface), precise enough compared to the typical angular variance (4 − 6 degree^2^) of beads attached to an inactivated motor in the magnetic tweezer.

Next, the PMF was restored by exciting the proteorhodopsin with 532 nm laser ^29-31^. Consequently the stators bound to the rotor one by one with the orientation angle of the magnetic bead increasing step by step, and the video recording was stopped until the angular change exceeded 30 degree. A typical experimental trace is shown in Fig. 1C. The stiffness was extracted from the Brownian motion trace *k* = *k*_*B*_*T*/< *δθ*^2^ >, where < *δθ*^2^ > was the angle variance when PMF was eliminated and the motor was inactivated, and the stall torque for each motor at each stator number was calculated from the motor resurrection trace.

### Precise calibration of the magnetic tweezer

A crucial issue was how to precisely calibrate the torsional stiffness of magnetic tweezer. Multiple effects, such as the bead incidentally attaching to somewhere other than the hook (e.g., the cell body), some stators still binding to the rotor, motion blur due to finite exposure time of the camera, effect due to finite frame rate of the camera, and stage drift-induced low frequency noise, might make the calibrated stiffness bigger or smaller ^32,33^. To eliminate the possibility of bead attaching to somewhere other than the hook, we perform further studies of the inactivated motors under no magnetic field, in which the rotor-hook-magnetic bead system approached free diffusive rotation, as shown in Fig. S1. If the bead was adhering to somewhere other than the hook, it is no longer free diffusion. If we calculated the stiffness with *k* = *k*_*B*_*T*/< *δθ*^2^ > assuming that the bead adhered to some linear constraint, the stiffness would be abnormally large up to ten thousands of pN×nm. To eliminate the possibility that some stators still binding to the rotor, we did the following. It was shown previously that the PMF was restored in less than 1 s upon 532 nm light illumination ^30^. If some stator still bound to the rotor, the motor would immediately resurrect with the direction of the magnetic bead changed once the 532 nm laser was on, otherwise the motor would resurrect after some period of time. Thus, we could get rid of this effect by judging whether the bead orientation immediately changed once laser was on. Evidently, there was no stators binding to the rotor before laser was turned on in Fig.1C. The camera frame rate was 300 frames/s and the exposure time was 3.3 ms. The trap relaxation time *t*_*relax*_ = *f*_*θ*_/*k* was typically about 74 ms where *f*_θ_ is the rotational frictional drag coefficient of the magnetic bead. The camera exposure time is much smaller than the trap relaxation time, so the motion blur was negligible ^33^. For a magnetic bead with rotation constrained by magnetic tweezer, the power spectrum for its angle trace is a Lorentzian *S*(*f*) = *A*/(1 + (*f*/*f*_*c*_)^2^) where *A* is a constant and *f*_*c*_ is the rolloff frequency (see Supporting Information). The camera frame rate (300 fps) was far greater than *f*_*c*_, which typically was about 2.6 Hz (Fig. 1E), so the effect of high frequency cutoff due to finite frame rate is negligible, and equivalently the effect on the variance of the bead position was negligible according to the Parseval theorem^32^. Low frequency drift of the sample stage would add low frequency noise to the power spectrum. To eliminate the effect of the low frequency drift, we simulated the Brown motion of bead trapped in magnetic tweezers with Langevin equation, then we filtered the trace using a high-pass filter over a range of cut-off frequencies from 0 to 0.3 Hz. We found that the variance of filtered angular position scaled linearly with the cut-off frequency as shown in Fig. S2. For our experimental data, the variance varied linearly with the cutoff frequency down to about 0.07 Hz, below which it was no longer linear due to drift-induced low frequency noise (Fig. S3). So we linearly fit the data between cutoff frequency of 0.07 to 0.3 Hz, and extrapolated it to 0 Hz to obtain the accurate variance of the bead angular position, as shown in Fig. S3.

### Two measurements at different magnetic strengths for individual motors

To further assure accuracy of the measurements of the stall torque, we performed two measurements at different magnetic strengths for each motor. The vertical distance between the magnets and the sample determines the magnetic strength for the tweezers. We selected two positions of the magnets, one of which corresponded to smaller strength *B*_*1*_ for the magnetic tweezers (position I) and the other corresponded to larger strength *B*_*2*_ (position II). At position II, the motion of the bead was recorded for about 360 s to calibrate the tweezer. The magnets were then moved to position I with the motion of the bead recorded for 360 s to calibrate the tweezer. Then the 532 nm laser was turned on to start motor resurrection. When motor resurrection proceeded long enough so that the angle change of the bead exceeded more than 30 degrees, the magnets were moved quickly to position II, reducing the angle change of the bead, as shown in Fig. 1C. We found that if the motor resurrection trace stepped stably, the relative difference for the two measurements usually satisfied:

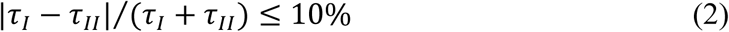

where *τ*_*I*_ = *k*_1_ < *θ*_1_ > and *τ*_*II*_ = *k*_2_ < *θ*_2_ > were the motor stall torque measured when the magnets were positioned at I and II immediately before and after position change, respectively, *k*_1_ and *k*_2_ represented the stiffness of the magnetic trap at position I and II, respectively, and < *θ*_1_ > and < *θ*_2_ > were the angle changes (relative to the original orientation of the magnetic bead) when the magnets were positioned at I and II immediately before and after position change, respectively. As the magnets were moved quickly from position I to position II (in less than 1 s), there was no change in stator number immediately before and after position change of the magnets. Therefore *τ*_*I*_ and *τ*_*II*_ are the measurements of the same motor stall torque at two different magnetic strengths, and should be equivalent. Equation (2) was a consistency check to make sure that errors in our measurements were within 10%. More examples of our experimental traces are shown in Fig. S4 and S5. Stall torques at different stator numbers were measured at magnets position I.

We measured resurrection traces for 20 motors. The average stall torques at different stator numbers are shown in Fig. 2, demonstrating a linear relationship, consistent with previous measurements at high loads ^4,15^. This also confirmed the linearity of the magnetic tweezer. We fit the data with a linear function and extracted the stall torque per stator to be 249.7 ± 37.4 *pN* · *nm*.

**Fig. 2.**
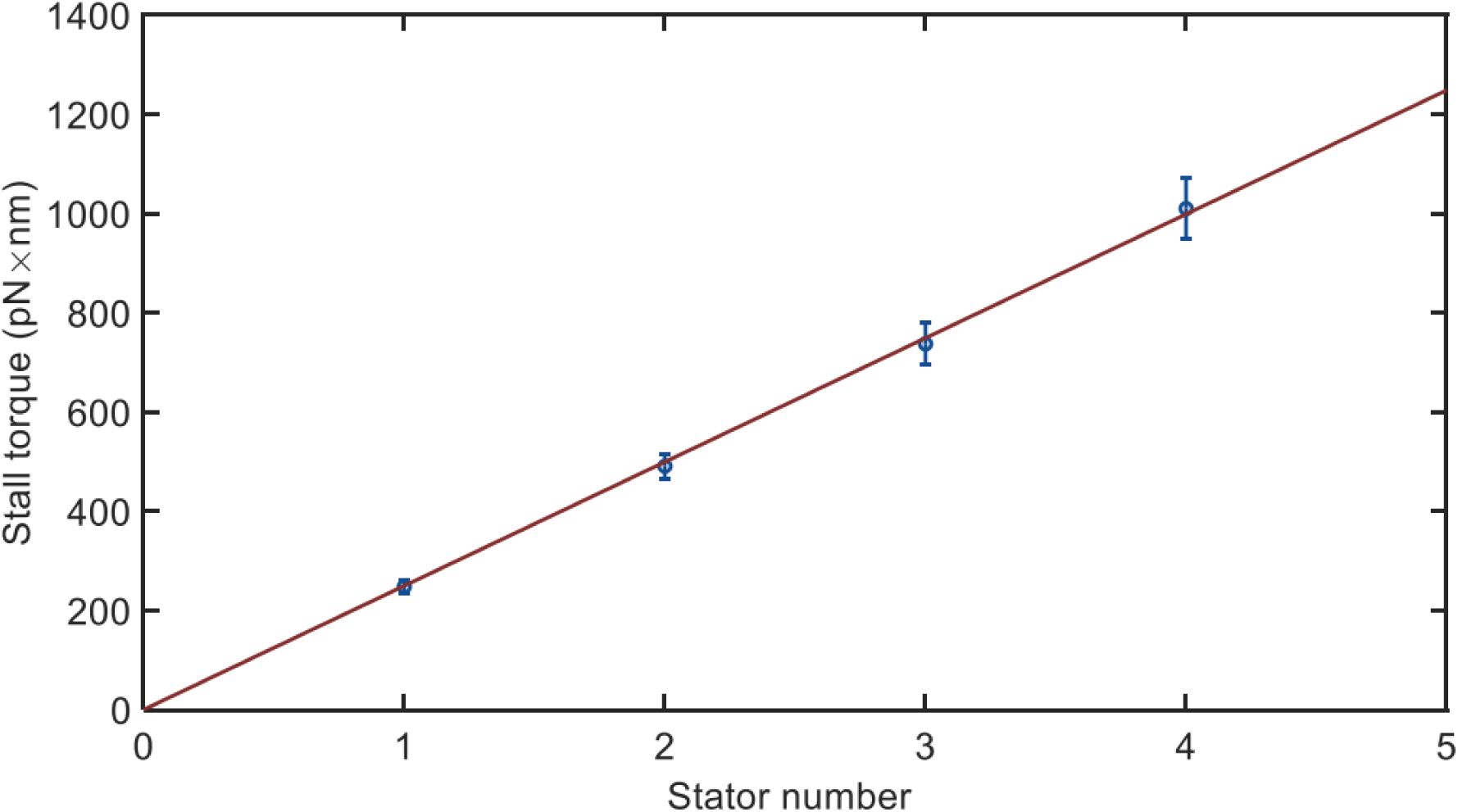
The stall torque as a function of stator number. The data were derived from 20 motor resurrection traces at stall. The red line was a linear fit the data. Error bars were the SEM.

Previous indirect measurements of the stall torque per stator were usually carried out with the bead assay by labeling 1.0-μm-diameter bead to the motor and assumed that the motor torque at this load is the same as the stall torque. To compare with previous measurements, we also performed motor resurrection experiments using a normal bead assay with 1.0-μm-diameter latex bead, obtaining an average value of 160.1 ± 13.6 *pN* · *nm* per stator for the motor torque at this load. Therefore our directly measured value of the motor stall torque was about 1.56 times the motor torque under load of 1.0-μm-diameter bead.

### Difference between the stall torque and the motor torque at high load

To explore the reason behind the difference between our measured stall torque and the motor torque under load of 1.0-μm-diameter beads, we sought to measure the torque-speed curve in the high-load region with the bead assay using different sizes of beads. Besides the plasmid pTrc99aPR, we transformed the strain JY9 with the plasmid pKAF131, which constitutively expresses sticky filament FliC^st^. We attached 0.75, 1.0, or 1.5-μm-diameter beads to shortened filament stubs of the motors, and carried out motor resurrection experiments using same 532 nm light condition as the tweezer experiments. Typical resurrection traces are shown in Fig. S6. We then constructed the torque-speed curves at different stator numbers at high load from the resurrection traces, as shown in Fig. 3. The motor torque at each stator number descends as the speed increases. This contributed to difference between our directly measured stall torque and the motor torque under load of 1.0-μm-diameter beads.

**Fig. 3.**
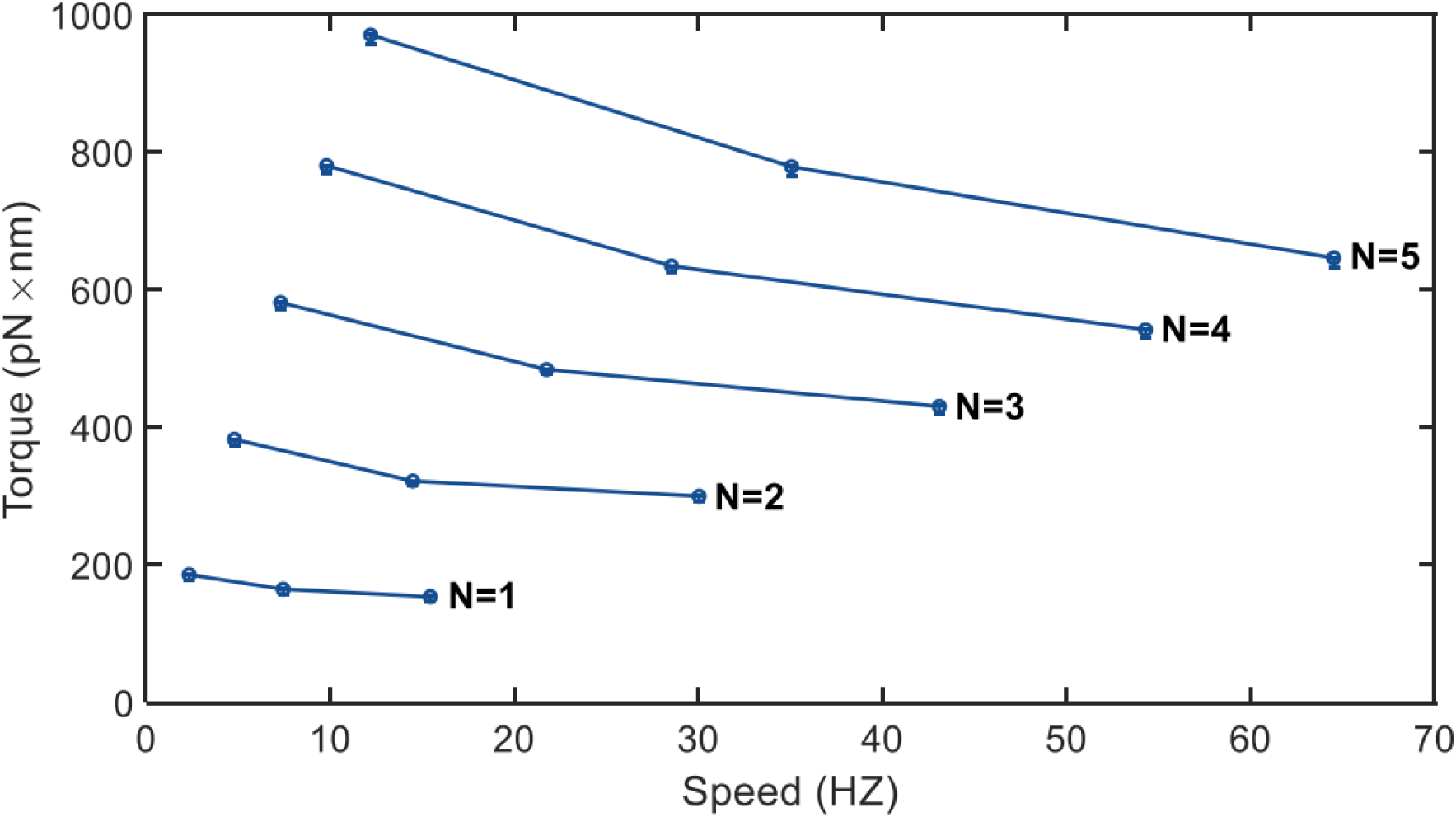
The torque-speed curve at different stator number in the high load region. From bottom to top, the stator number for each line was from 1 to 5. Data were from the motor resurrection experiments under loads of 1.5, 1.0, and 0.75-μm-diameter beads. The numbers of motors observed at each load were 32, 42, and 23, respectively. Error bars were the SEM.

Another contribution came from the calculation of the rotational viscous drag coefficient of the load in the bead assay. The motor torque in the bead assay was calculated by multiplying this drag coefficient with the motor speed. Usually in calculating the drag coefficient, the filament stub and the bead was assumed to be rotating in an infinitely large environment, neglecting the hydrodynamic surface effect from the cell body. As the drag coefficient came mostly from rotation of the bead, we sought to estimate the hydrodynamic surface effect on rotation of the bead. When the surface effect was neglected, the rotational drag coefficient of the bead is

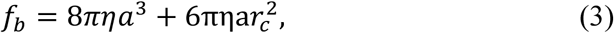

where *η* is the viscosity of the medium, *a* is the radius of the bead, and *r*_*c*_ is rotational radius of the bead. From our experiments, the rotational radius *r*_*c*_ was about 210 nm. For simplicity we treated the upper surface of the cell body as an infinite plane parallel to the sample glass coverslip, and neglected the hydrodynamic surface effect from the glass coverslip. The rotational plane of the bead was usually not parallel to the cell body surface, with an average intersection angle of about 50 degree. On average the bead stuck to the filament at the length of about 1000 nm, so the distance *s* from the center of the bead to the cell body surface was approximately 750 nm. Motion of the bead could be decomposed into two directions parallel and perpendicular to the plane of cell body. So the actual rotational drag coefficient of the bead including the surface effect is ^34^:

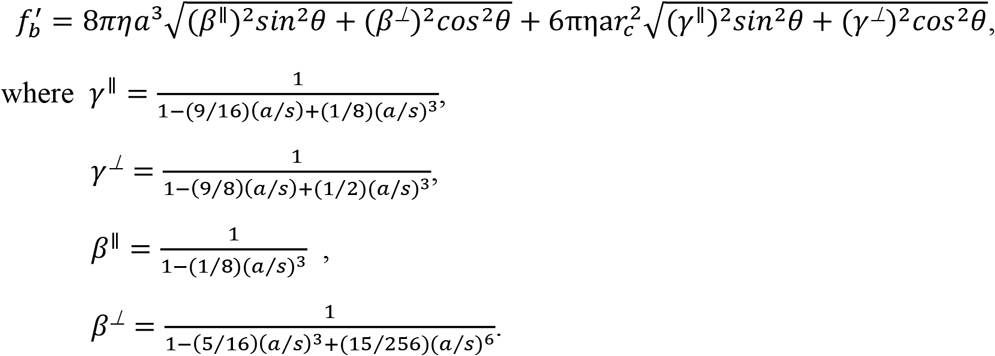

So 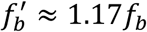. The actual rotational drag coefficient was 1.17 times that without the surface effect as was usually done.

According to trend of the torque-speed curve in the high load region (Fig. 3), the stall torque was about 1.22∼1.32 times the motor torque under load of 1.0-μm-diameter beads. Combining with the factor of about 1.17 from the surface effect, the stall torque was about 1.42∼1.54 times the motor torque under load of 1.0-μm-diameter beads. This explains the difference between our directly measured stall torque and the motor torque at hight load.

### Summary and Discussion

In past several decades, magnetic tweezers were widely used for studying nucleic acid enzymes ^35-37^. The torsional stiffness ranged from several thousands of *pN* · *nm*/rad to ten thousands of *PN* · *nm*/*rad*, close to the magnitude of the stall torque of the flagellar motor. Recently several works have applied magnetic tweezers to study the bacterial flagellar motor, finding that the magnetic field does no harm to the motor ^28,38^. In this work, we took advantage of the magnetic tweezer to quantitatively measure the stall torque of the flagellar motor, which was an important parameter for modelling the motor and further understanding the working mechanism of motor.

Here, we performed careful calibration of the magnetic tweezer by ruling out multiple possible effects. We then applied magnetic tweezer to directly measure the stall torque per stator. We made measurements of the stall torque at two magnetic strengths for each motor to verify the accuracy of our measurements. The stall torques we measured are proportional to the stator number, further confirming the linearity of the magnetic tweezer. Our directly measured stall torque is about 1.52 times the motor torque under load of 1.0-μm-diameter beads which was previously taken to be the stall torque in indirect measurements. We explained the difference by measuring the shape of the torque-speed curve in the high-load region and by estimating the hydrodynamic surface effect from the cell body on rotation of the bead.

We sought to estimate the number of protons *n* each stator passes per revolution. When the motor rotates infinitely slowly, the motor efficiency is 1, namely,

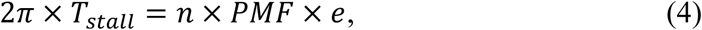

where *T*_*stall*_ is the stall torque per stator, and *e* is the proton charge. The PMF in a wild-type *E. coli* K12 cell was about 190 mV ^39^. In the current study, the PMF established by light-driven proteorhodopsin proton pump is slightly smaller. As the motor speed varies linearly with the PMF ^40^, we can use the ratio of the motor speeds for motors driven with wildtype PMF and motors driven with proteorhodospin-pumped PMF to extract to latter PMF. We compared the ratio at each stator number using the motor resurrection data in this study and the wildtype motor resurrection data ^8^, both under load of 1.0-μm-diameter bead, as shown in Fig. S7. The ratio is 1.12. So the proteorhodopsin-pumped PMF in our experiments is about 170 mV. If we used motor torque under load of 1.0-μm-diameter bead measured above (160.1 *pN* · *nm*) as the stall torque as was previously normally assumed, the number of protons each stator passes per revolution would be 37, consistent with the previous measurement ^22^. This further confirmed that our estimate of the proteorhodopsin-pumped PMF was correct.

Therefore, each stator passes 58 ± 9 protons per revolution, or equivalently, each stator passes 2.23 ± 0.34 protons per step as the motor takes 26 discrete steps per revolution ^23^. This is consistent with the findings that each stator takes two “power strokes” per step and each power stroke is induced by one proton passing through one of the two proton channels in a stator ^2,41,42^. This supported the model that the motor rotation was tightly coupled with the proton transport ^26^.

## Materials and Method

### Strains and plasmids

All strains for this study are derivatives of *E. coli* K12 strain RP437. JY9 (Δ*cheY fliC*) carries a mutated gene *flgE* on chromosome that expresses the protein with a tetracysteine motif CCXXCC at codon 220. The plasmid pTr99aPR expresses proteorhodopsin under control of an IPTG-inducible promoter. The plasmid pKAF131 expresses the sticky flagellar filaments to readily adsorb polystyrene beads for the bead assay.

### Optics

We constructed a system combing optical trap, magnetic tweezer, 532 nm laser illumination, and bright-field imaging based on a Nikon Ti-U inverted microscope. A scheme of the set-up is shown in Fig. 1B. The optical trap was constructed with a 1064 nm laser beam (AFL-1064-33-B-FA; Amonics), which was expanded 5 times with two convex lens, reflected by a dichroic mirror (ZT1064rdc, Chroma), and focused into a diffraction-limited spot with a water-immersion objective (Nikon Plan Apo vc 60×/1.20 WI). A 532 nm fiber-coupled laser light (MGL-III-532, Cnilaser) was expanded 13 times, and focused onto the back focal plane of condenser lens by a long-focus lens. The light was reflected by a dichroic mirror (ZT543rdc-UF2, Chroma) between the condenser lens and the long-focus lens, and expanded into a parallel beam by the condenser lens. There was a 1.5-mm-diameter center openning in the holder of magnetic tweezer that allowed passage of the 532 nm laser and the bright-field illumination light. The holder was placed on a 3D PC-controlled motorized platform (MTS202, BeiJing Optical Century Instrument Co.,LTD) between the condenser and the sample. The density of 532 nm light for motor resurrection was 3.8 mW/mm^2^, at which the effect of the proteorhodopsin was saturated (Fig. S8). The light for bright-field microscopy was provided by a Halogen lamp illuminating the sample from above. All convex lenses were from Thorlabs.

### Labeling of magnetic beads and latex beads

Cells were grown in 3 ml of T-broth with 100 μg/ml ampicillin at 33 °C to optical density (OD_600_) of 0.4, then 3 μl all-trans-retinal and 3 μl of 20 mM Maleimide-PEG2-Biotion (MPEGB) ^29^ were added, and cells were re-cultivated for about 1 hour until the OD_600_ reached 0.5-0.6. Cells were harvested by washing twice with motility buffer (10 mM potassium phosphate, 0.1 mM EDTA, 10 mM lactate, and 70 mM NaCl at pH 7.0). They were mixed with 3 μl of 20 mM MPEGB at 30 °C for 1 hour with shaking to biotinylate the hooks, re-washed with motility buffer twice, and ultimately re-suspended in 300 μl motility buffer for subsequent resurrection experiments. The sample chamber was constructed by using two layers of double-sided sticky tape as a spacer between a glass slide and a glass coverslip coated with poly-L-lysine, then was placed in a baker oven at 70 °C for about 10 minutes and subsequently allowed to cool down. Cells were flown into the chamber and allowed to stick onto the coverslip in 7 minutes. Unstuck cells were washed away with motility buffer containing 23 mM NaN_3_, then solution of 2.8-μm-diameter streptavidin-coated magnetic beads (11205D, Thermo Fisher) was added into the chamber to attach to the biotinylated hook spontaneously. Unattached magnetic beads were washed away, and then solution of 0.0015% w/v 1.0-μm-diameter biotin-labeled latex beads (F8768, Thermo Fisher) was drawn slowly into the chamber. The chamber was then sealed with Apiezon vacuum grease. The shutter for the 1,064 nm laser was opened to capture a latex bead, and the sample stage was then translated so that a straptavidin-coated magnetic bead attached to a motor was moved close to and stuck to the captured latex bead. The 1064 nm light for the optical trap was immediately shut off. The beads were observed with bight-field microscopy, a region of interest (ROI) was chosen to cover the magnetic bead and the latex bead, and images and videos were recorded using a CMOS camera (Thorlabs, DCC1545M).

### Data analysis

Data analysis was carried out using custom scripts in MATLAB. A latex bead was stuck to the magnetic bead as a fiducial marker to indicate its orientation, as sketched in the inset in Fig. 1B. An example bright-field image of the two beads is shown in Fig. 1A. The focusing plane was chosen so that the latex bead was in focus and usually the magnetic bead was slightly out of focus. To calculate the angle accurately, we adapted the algorithms described in previous studies ^36,43^ (see **Algorithm for angle detection**). Determination of angle steps in motor resurrection at stall was carried out by using a step-finding algorithm described previously ^8^. For the bead assay, the motor torque at different high loads was computed with the formula *T*_*motor*_ = (*f*_*b*_ + *f*_*f*_) ×*ω*, where *f*_*b*_, *f*_*f*_ were the rotational drag coefficients of the bead and the filament stub, respectively, and ω was the rotational speed of the motor ^21^.

### Algorithm for angle detection

The reference image that displays similar ring patterns as the magnetic bead image is 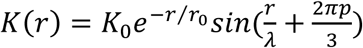, where *K*_0_ is a constant, *r* is the distance from the image center, *r*_0_ is a decay length, *λ* is the fringe spacing, and *p* determines the shift in the ring pattern ^43^. The values of the parameters we used were *K*_0_ = 1, *r*_0_ = 30, *λ* = 4, *p* = 0.5. The reference image was convoluted with the real image to find the center of the magnetic bead at single pixel precision (83 nm). To further get sub-pixel resolution, each pixel of the real image was divided into 5×5 sub-pixels, the intensities of which were obtained by linear interpolation. Then autocorrelation calculation was performed with a shift grid of 11×11 around this center of the magnetic bead to get the center position at sub-pixel precision. A ring-shaped region (inner and out radiuses are 15 and 35 pixels, respectively) was selected from the real image around the center of magnetic bead that covered the image of the latex bead. It was then transformed into a ploar intensity profile with linear interpolation ^43^, using a polar coordinate centered on the magnetic bead with angular coordinate segmented into steps of 0.2°. The polar profile was summarized over radius *r* to obtain the angular profile: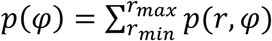. Then *p*(*φ*) was cross correlated with *p*^−1^(*φ*), which was derived from the mirror image of the ring-shaped image about the *x*-axis, to derive the shift *φ*_0_ at highest correlation, and the orientation of the magnetic bead was 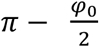.

## Acknowledgements

This work was supported by National Natural Science Foundation of China Grants (11925406, 21573214, and 11872358), a grant from the Ministry of science and technology of China (2016YFA0500700), and a grant from Collaborative Innovation Program of Hefei Science Center, CAS (2019HSC-CIP004). B.W. is supported by the National Postdoctoral Program for Innovative Talents (BX2018211928).

## Author Contributions

J.Y. and R.Z. planned the work; B.W. and G.Y. performed the measurements; J.Y., R.Z., and B.W. wrote the paper; B.W. and G.Y. contributed equally to this work.

