## Supporting information for "Direct precise measurement of the stall torque of the flagellar motor in *E. coli*"

### Power spectrum for the rotation of the magnetic bead in magnetic tweezer

For a magnetic bead attached to an inactivated motor in magnetic tweezer, the Langevin equation for its rotation is

$$f_\theta \frac{d\theta(t)}{dt} + k\theta(t) = F_{therm}(t), \tag{1}$$

where $f_\theta$ is the rotational drag coefficient of the bead, $k$ is the torsional stiffness of the magnetic tweezer, and $F_{therm}(t)$ was the random force due to Brown collision, with $< F_{therm}(t) > = 0$, $< F_{therm}(t)F_{therm}(t+\tau) > \sim\delta(\tau)$. In equation (1), $t$ was replaced by $t+\tau$, and then equation was multiplied by $\theta(t)$, subsequently integrated for $t$ over 0 to $T$ at both sides.

$$f_\theta \frac{1}{T}\frac{d\int_0^T \theta(t)\theta(t+\tau)dt}{d\tau} + k\frac{1}{T}\int_0^T \theta(t)\theta(t+\tau)dt = \frac{1}{T}\int_0^T \theta(t)F_{therm}(t+\tau)dt \tag{2}$$

For $T \to \infty$ and $\tau > 0$, equation (2) was transformed into:

$$f_\theta \frac{dCorr(\tau)}{d\tau} + kCorr(\tau) = 0, \tag{3}$$

where $\mathrm{Corr}(\tau) = < \theta(t)\theta(t+\tau) > = \lim_{T\to\infty}\frac{1}{T}\int_0^T \theta(t)\theta(t+\tau)dt$ is the temporal correlation function. So the temporal correlation function had a form of exponential decay:

$$Corr(\tau) = Corr(0)\exp(-\frac{k}{f_\theta}\tau) \tag{4}$$

The associated power spectrum was then obtained according to the Wiener-Khinchin theorem to be

$$S(f) = A/(1+(f/f_c)^2), \tag{5}$$

a Lorentzian where $A$ was a constant, and $f_c$ was the rolloff frequency with $2\pi f_c = k/f_\theta$.

An example of the power spectrum for the trace of a magnetic bead attached to an inactivated motor in magnetic tweezer (Fig. 1C) is shown in Fig. 1E. The Lorentzian fit gave $f_c$ =2.65 Hz. The stiffness was 3033.7 $PN \cdot nm/rad$ from $k = k_BT/< \delta\theta^2 >$. So the drag coefficient was 182.2 $PN \cdot nm \cdot s$, which was consistent with that estimated from hydrodynamic calculate on.

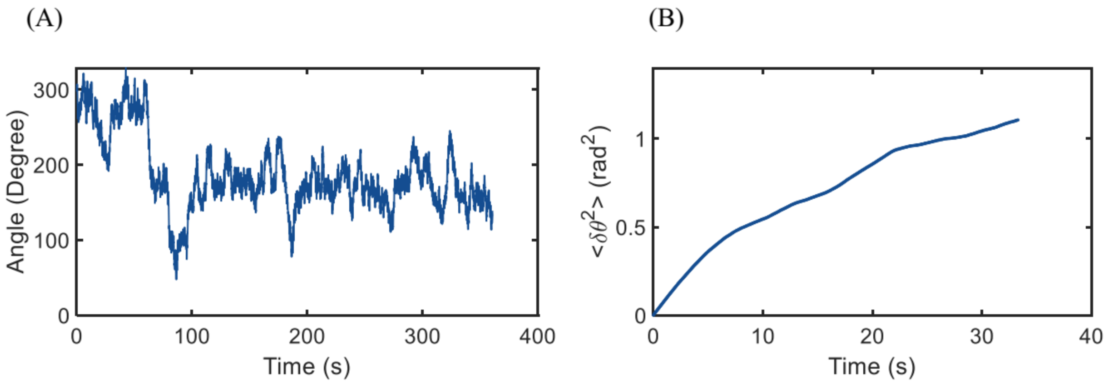


Fig. S1. The free diffusion of the magnetic bead attached to an inactivated motor with no magnetic field. (A) The orientation of the bead as a function of time. (B) Mean-squared angle displacement as a function of time for the trace in (A). The drag coefficient was calculated to be about 241.7 $PN \cdot nm \cdot s$ from the diffusion trace.

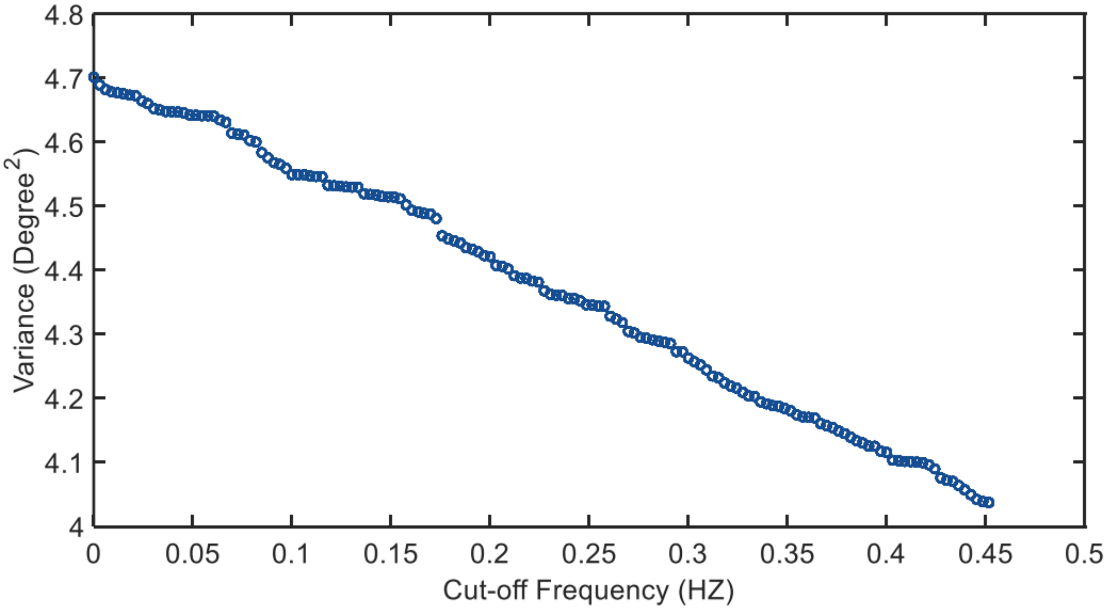


Fig. S2. Angular variance of a simulated trace (high-pass filtered) of a magnetic bead constrained by magnetic tweezer, as a function of cut-off frequency of the high-pass filter.

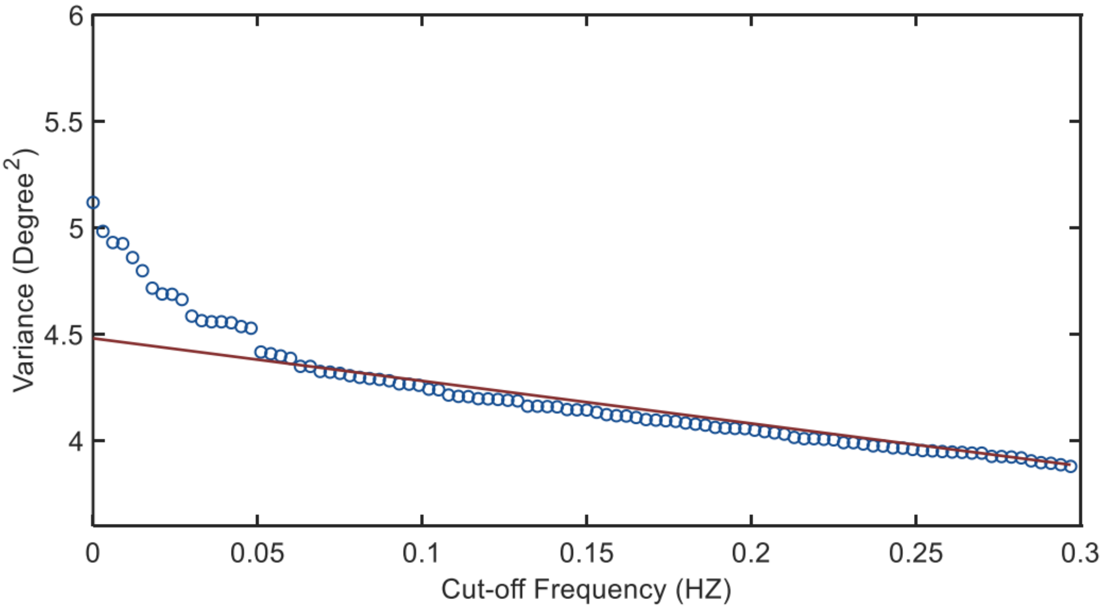


Fig. S3. Angular variance of an experimental trace of a magnetic bead attached to an inactivated motor in magnetic tweezer. The trace was high-pass filtered, and the angular variance was plotted as a function of the cut-off frequency. The red line was a linear fit to the data points above 0.07 Hz to obtain the unbiased variance by extrapolating to 0 Hz.

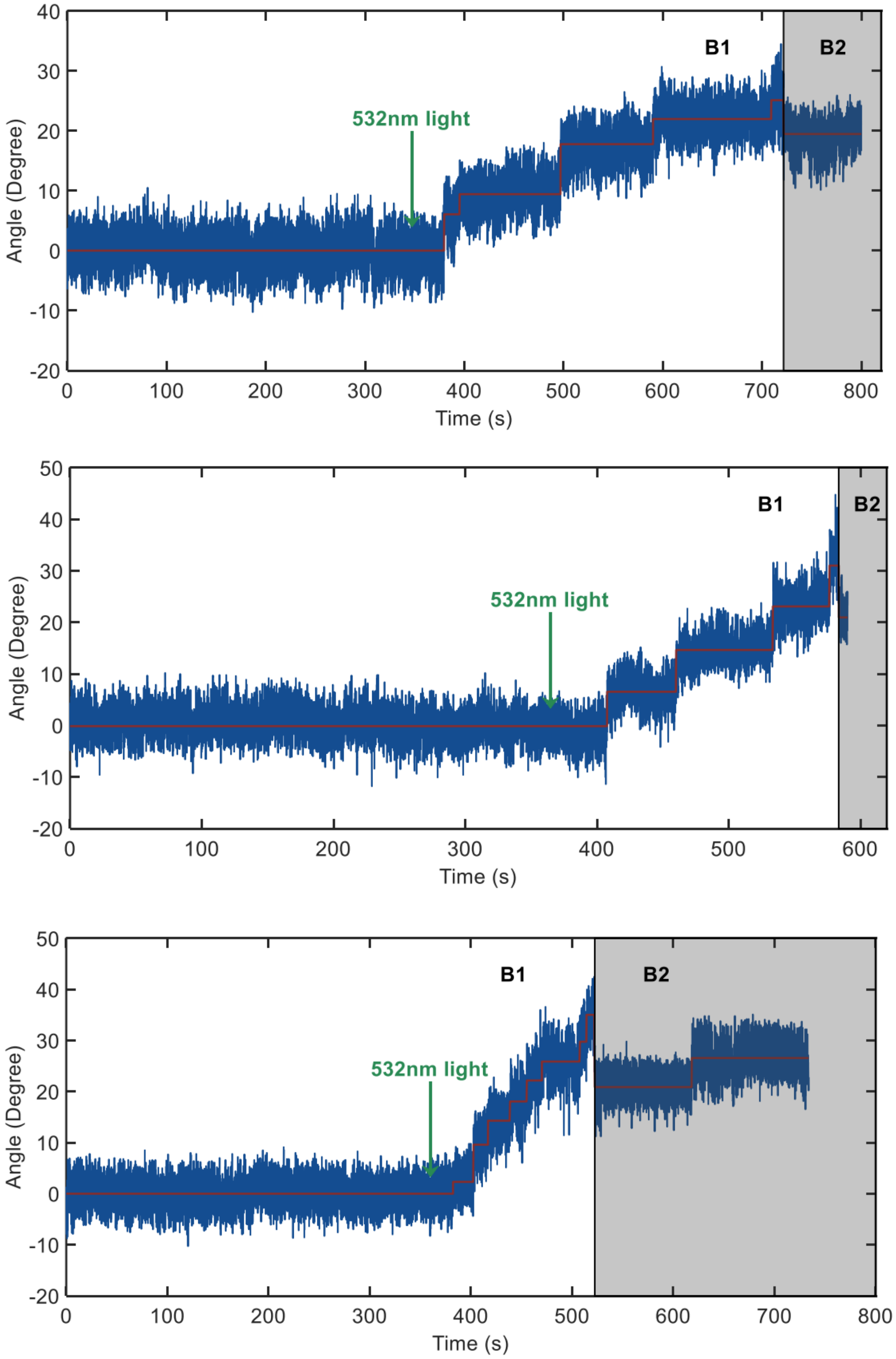


Fig. S4. More examples of the traces of motor resurrection at stall.

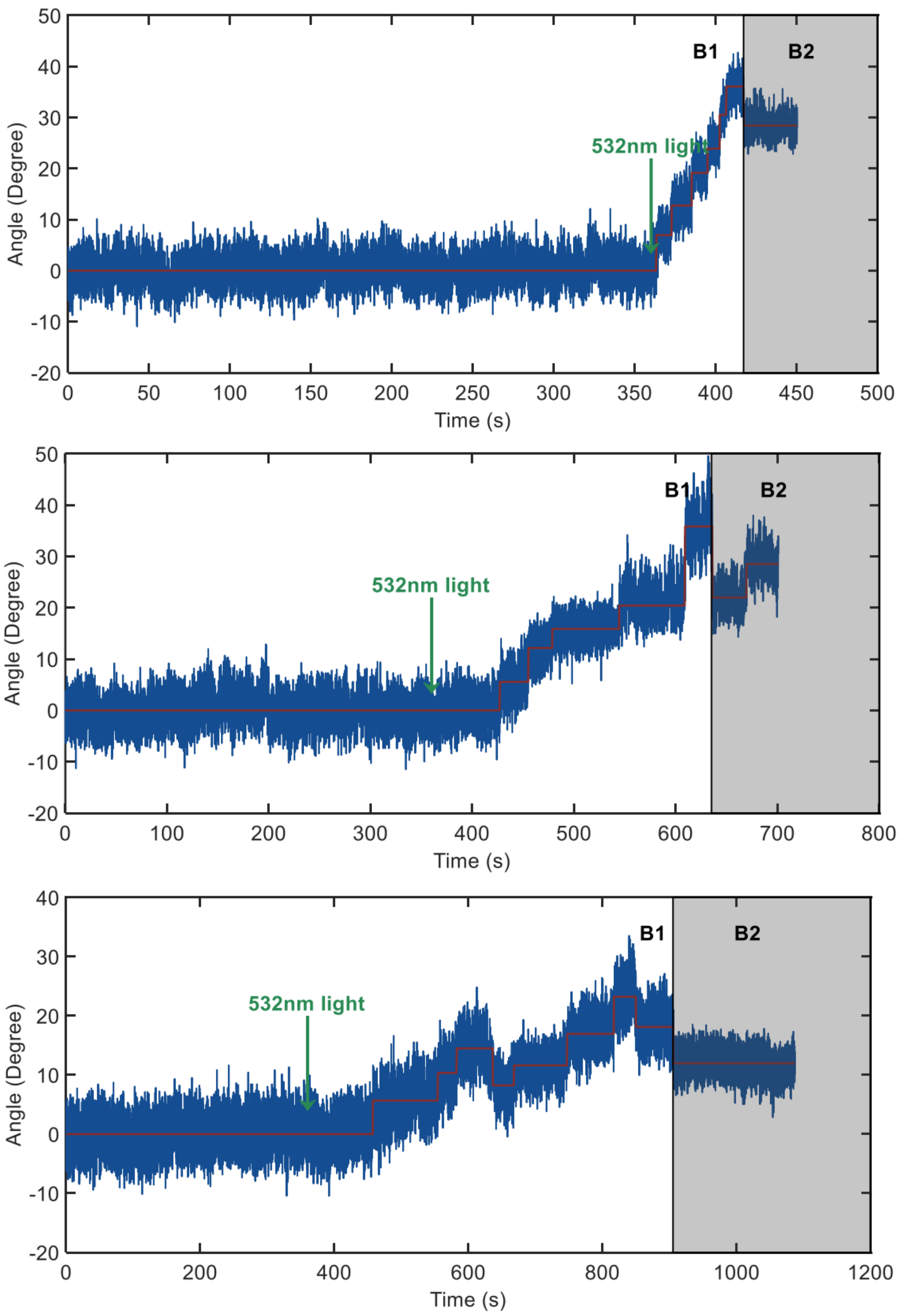


Fig. S5. More examples of the traces of motor resurrection at stall.

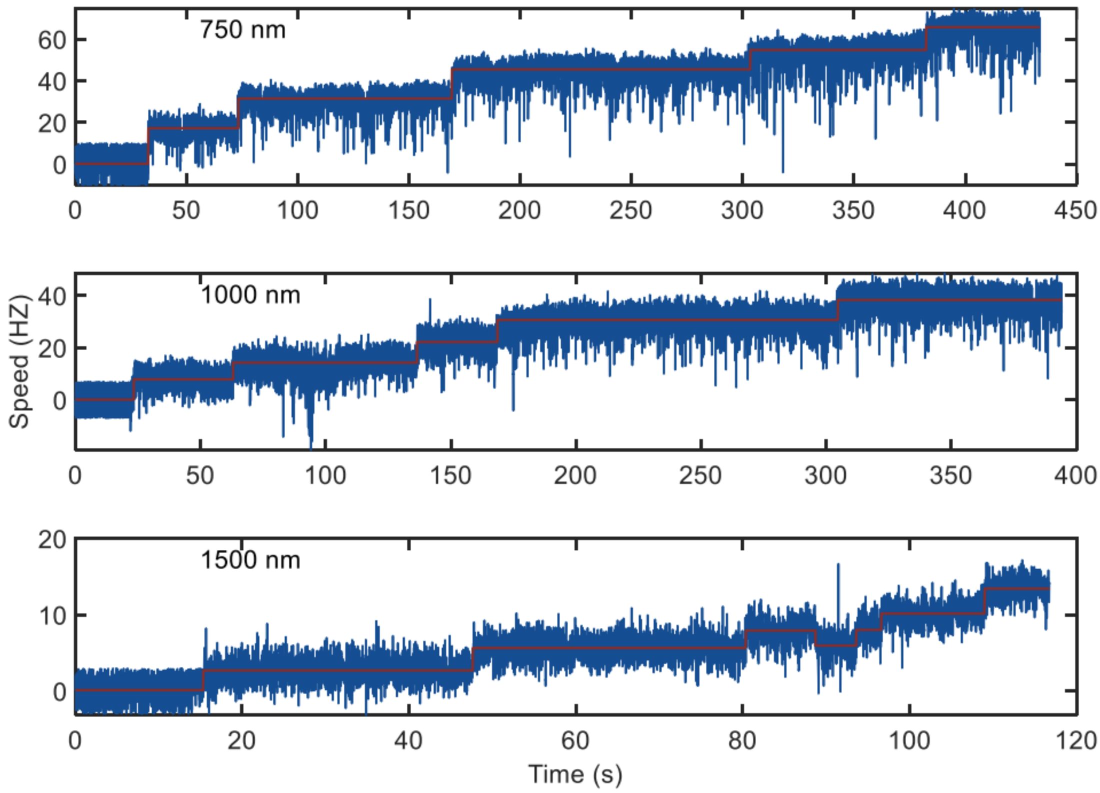


Fig. S6. Typical resurrection traces for motors labelled with 0.75, 1.0, and 1.50-μm-diameter beads (from top to bottom panels). The red lines were the speed steps identified by step-finding algorithm.

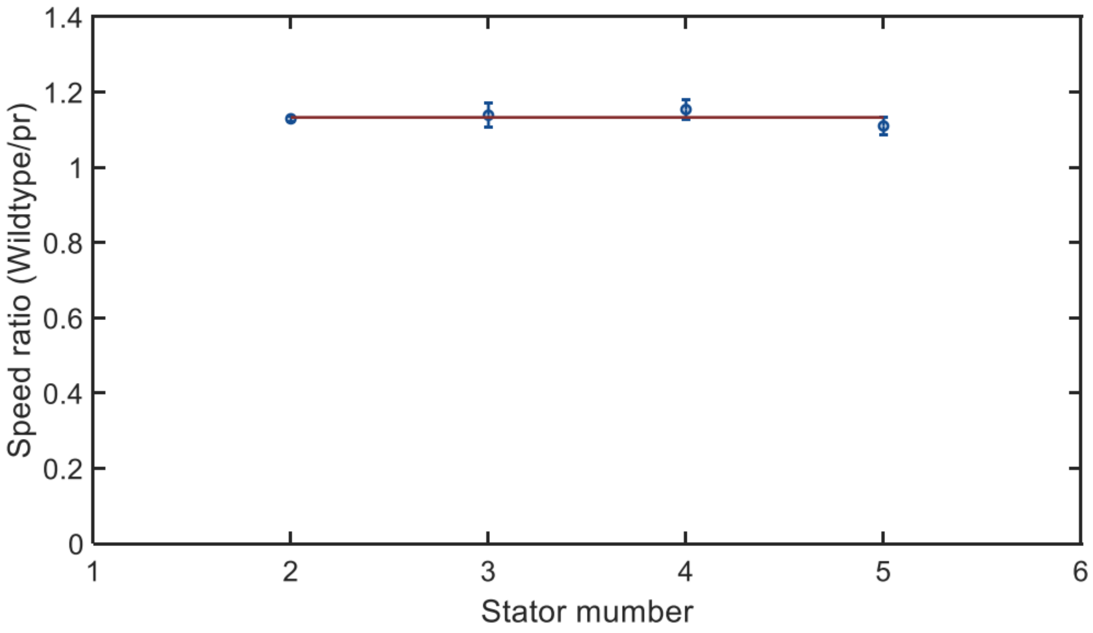


Fig. S7. The ratio of motor speeds at each stator number for wildtype cells and cells with PMF established by light-driven proteorhodopsin. The value of the red line was 1.12 by fitting the data with a constant.

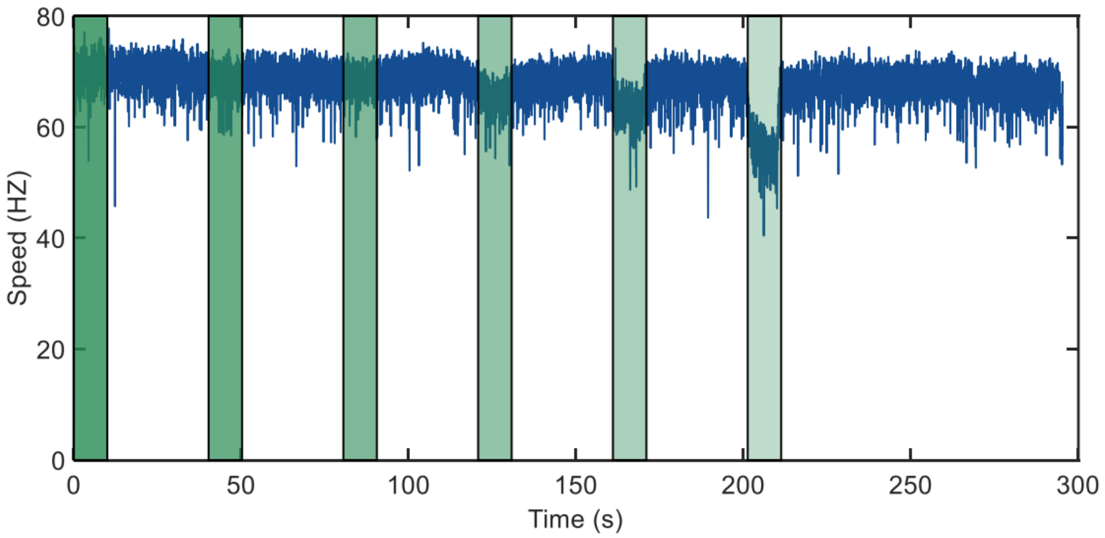


Fig. S8. Motor speed at various intensities of the 532 nm laser. The light densities in the green zones ranging from dark to light were 3.8, 3.1, 2.3, 1.5, 0.76, and 0.38 $mW/mm^2$. The light density of the other white sections was 3.8 $mW/mm^2$. Red line was the average value of motor speeds in each section.
